# GenMPI: Cluster Scalable Variant Calling for Short/Long Reads Sequencing Data

**DOI:** 10.1101/2022.04.01.486779

**Authors:** Tanveer Ahmad, Joseph Schuchart, Zaid Al Ars, Christoph Niethammer, José Gracia, H. Peter Hofstee

## Abstract

Rapid technological advancements in sequencing technologies allow producing cost effective and high volume sequencing data. Processing this data for real-time clinical diagnosis is potentially time-consuming if done on a single computing node. This work presents a complete variant calling workflow, implemented using the Message Passing Interface (MPI) to leverage the benefits of high bandwidth interconnects. This solution (GenMPI) is portable and flexible, meaning it can be deployed to any private or public cluster/cloud infrastructure. Any alignment or variant calling application can be used with minimal adaptation. To achieve high performance, compressed input data can be streamed in parallel to alignment applications while uncompressed data can use internal file seek functionality to eliminate the bottleneck of streaming input data from a single node. Alignment output can be directly stored in multiple chromosome-specific SAM files or a single SAM file. After alignment, a distributed queue using MPI RMA (Remote Memory Access) atomic operations is created for sorting, indexing, marking of duplicates (if necessary) and variant calling applications. We ensure the accuracy of variants as compared to the original single node methods. We also show that for 300x coverage data, alignment scales almost linearly up to 64 nodes (8192 CPU cores). Overall, this work outperforms existing big data based workflows by a factor of two and is almost 20% faster than other MPI-based implementations for alignment without any extra memory overheads. Sorting, indexing, duplicate removal and variant calling is also scalable up to 8 nodes cluster. For pair-end short-reads (Illumina) data, we integrated the BWA-MEM aligner and three variant callers (GATK HaplotypeCaller, DeepVariant and Octopus), while for long-reads data, we integrated the Minimap2 aligner and three different variant callers (DeepVariant, DeepVariant with WhatsHap for phasing (PacBio) and Clair3 (ONT)). All codes and scripts are available at: https://github.com/abs-tudelft/gen-mpi

## 1 INTRODUCTION

Next Generation Sequencing (NGS) technologies can produce high throughput, less erroneous and higher depth/coverage sequencing data at low costs. These high-throughput sequencing technologies are making their way from research to the field in a wide range of applications ranging from clinical diagnostics to agriculture. Depending on the experiment design type, the need of sequencing coverage varies. A typical 300x coverage human genome dataset size exceeds 2.3 TBytes (Zook et al., 2016). Processing such large datasets on a single-node can take up to several days. For a number of applications employing higher coverage sequencing data and/or requiring urgent sequencing results, this exceeds acceptable time constraints.

Sequencing coverage influences both accuracy and the sensitivity of genomics analysis. In pediatric brain-tumor studies (Kline et al., 2016), more than 200x coverage for the tumor sample and more than 100x coverage for the normal sample are collected for better precision for the tumors of concern. Recent studies show that whole exome sequencing (WES)/whole genome sequencing (WGS) helps with diagnosis, decision making, and treatment of fetal diseases (Becher et al., 2020). Although WES is normally used to detect fetus anomalies in prenatal and perinatal testing, which only targets protein-coding regions of genes in a genome, this type of sequencing needs urgent and fast sequencing analysis due to the time-critical nature of these tests. Sequencing can also enable finding some rare hereditary disorders and genetic variants associated with specific diseases in newborn screening. A recent study that uses genomic sequencing for newborn screening (Roman et al., 2020) showed that some of enrolled healthy newborns and children with metabolic diseases or hearing loss exhibited pathogenic variants associated with hereditary breast or ovarian cancer and a pathogenic variant in the gene associated with Lowe syndrome. This shows that sequencing in newborn screening can play a vital role in timely diagnosis and treatment of diseases and ultimately will lead to urgent need of processing genomics data in such time-critical clinical settings.

Many consortia, associations and government disease diagnosis and drug regulatory agencies stipulate guidelines/protocols for genomics sequencing as a standard tool in diagnosis and treatment of some fatal diseases (Allegretti et al., 2018). Many medical and diagnosis centers, particularly in developed countries, have started sequencing as a regular practice for prenatal and perinatal testing, newborn screening, genetic and cancerous disease diagnostic and personalized treatments. In the coming decades, as sequencing becomes a normal practice for human health and other types of research, locally available compute infrastructure to any organization will not be adequate to fulfill the sequencing requirements. At the same time, large computing infrastructure also requires human resources and incurs power and maintenance costs.

Mostly, genome sequencing machines produce FASTQ format raw data for a given sample (Deorowicz and Grabowski, 2011). Mapping this raw data to a reference genome, sorting the resultant Sequence Alignment/Map (SAM) reads according to chromosomes and their positions, and finally removing the duplicate reads are some of the standard practices before actual variant detection. A large number of tools is available for the aforementioned methods but almost all are developed for use in a single compute node context. Scaling these tools for cluster environments for both scalability and reproducibility is an ongoing challenge. In the past decade, many cluster-scaled solutions have been created, almost all these solutions use big data frameworks like Apache Spark (Apache, 2019b) or Apache Hadoop (Apache, 2019a; Dean and Ghemawat, 2008) for distributing and work scheduling. Data formats like Apache Parquet, Apache Arrow, Apache Avro have been explored extensively in conjunction with these frameworks to store and process genomic data efficiently. These frameworks include ADAM (Massie et al., 2013), SparkGA2 (Mushtaq et al., 2019), VC@Scale (Ahmad et al., 2021) and Halvade (Decap et al., 2015). Due to many underlying dependencies, inefficient memory usage, issues related to scalability, cluster deployment challenges as well as incompatible data formats, solutions based on these frameworks are still not widely used in the mainstream Bioinformatics community. While the Message Passing Interface (MPI) has been used previously for parallelization of genomic algorithms on HPC systems like pBWA (X et al., 2012), in this work we explore a comprehensive approach towards using MPI for cluster scale parallelization of genomics applications and complete variant calling workflows.

We expect to benefit from the following advantages that MPI has over traditional big data frameworks for genomics workflows.

- Bare-metal performance and linear scalability of existing applications;
- Portable access to low-level network capabilities;
- Little to no extra memory overheads otherwise incurred in big data frameworks;
- Efficient MPI I/O performance on parallel file systems (like Lustre (Lustre, 2020), GPFS (IBM, 2020)).

In the following, we list the main contributions of this work.

- MPI-based parallelization of BWA-MEM (Li and Durbin, 2009) and Minimap2 (Li, 2016), compressed input FASTQ files can be provided as separate files to parallelize streaming of input while reading of uncompressed FASTQ will be parallelized internally. SAM output is tested on both POSIX and shared MPI I/O.
- Sorting, indexing and duplicate reads removal (if necessary) can be performed through a queue employing low-level network atomic operations (if input is already chunked based on chromosomes) or through MPI based bitonic sorting.
- For short reads, GATK HaplotypeCaller (Institute, 2010), Octopus (Cooke et al., 2021), and DeepVariant (Poplin et al., 2018) are used in combination with the MPI RMA-based queue for parallel chromosomes processing on a cluster.
- For long reads, DeepVariant, DeepVariant with WhatsHap (Martin et al., 2016) for phasing, and Clair3 (Zheng et al., 2021) variant callers are used in combination with the MPI RMA-based queue for parallel chromosome processing on a cluster.
- Resultant VCFs are merged through Bcftools (Danecek et al., 2021) to generate a single combined VCF (variant calling file) and to insure variants correctness.
- SNP/INDEL accuracy/precision/F1 tests are performed through hap.py (Krusche, 2021) against GIAB v4.2.1 (GIAB, 2021) benchmark set for the HG002 dataset.

The rest of this article is organized as follows. In Section 2 we describe our implemented methods in detail for both pre-processing and variant calling for short and long reads NGS data, followed by Section 3 where we compare the methods integrated into this approach with the existing workflows for both performance, accuracy, and scalability. In Section 4, run-time, accuracy, scalability, portability, and cost efficiency are discussed briefly. Finally we conclude this work in Section 5.

## 2 METHODS

We have constructed both short and long reads based cluster scalable variant calling workflows. For pre-processing of short and long reads NGS data, the *BWA-MEM* (Li and Durbin, 2009) aligner with sorting (*Samtools* (Li, 2009)) and mark duplicate (*Sambamba* (Tarasov et al., 2015)) are used for the former, while *Minimap2* (Li, 2016) with sorting (*Samtools*) is used for the latter. As shown in Figure 1, for short reads we use three different variant callers like *Octopus* and *DeepVariant* as well as GATK best practices variant calling pipeline using *HaplotypeCaller*. Similarly, for PacBio long reads, DeepVariant and DeepVariant with chromosomes phasing using *WhatsHap* have been used while *Clair3* variant caller is used for Oxford Nanopore Technologies (ONT) data as recommended by ONT (medaka, 2021). These workflows can be run through both cluster workload managers like Slurm (Slurm, 2020) and PBS (OpenPBS, 2020). The following sections provide a more detailed description of both short and long reads workflows implementation.

**Figure 1.**
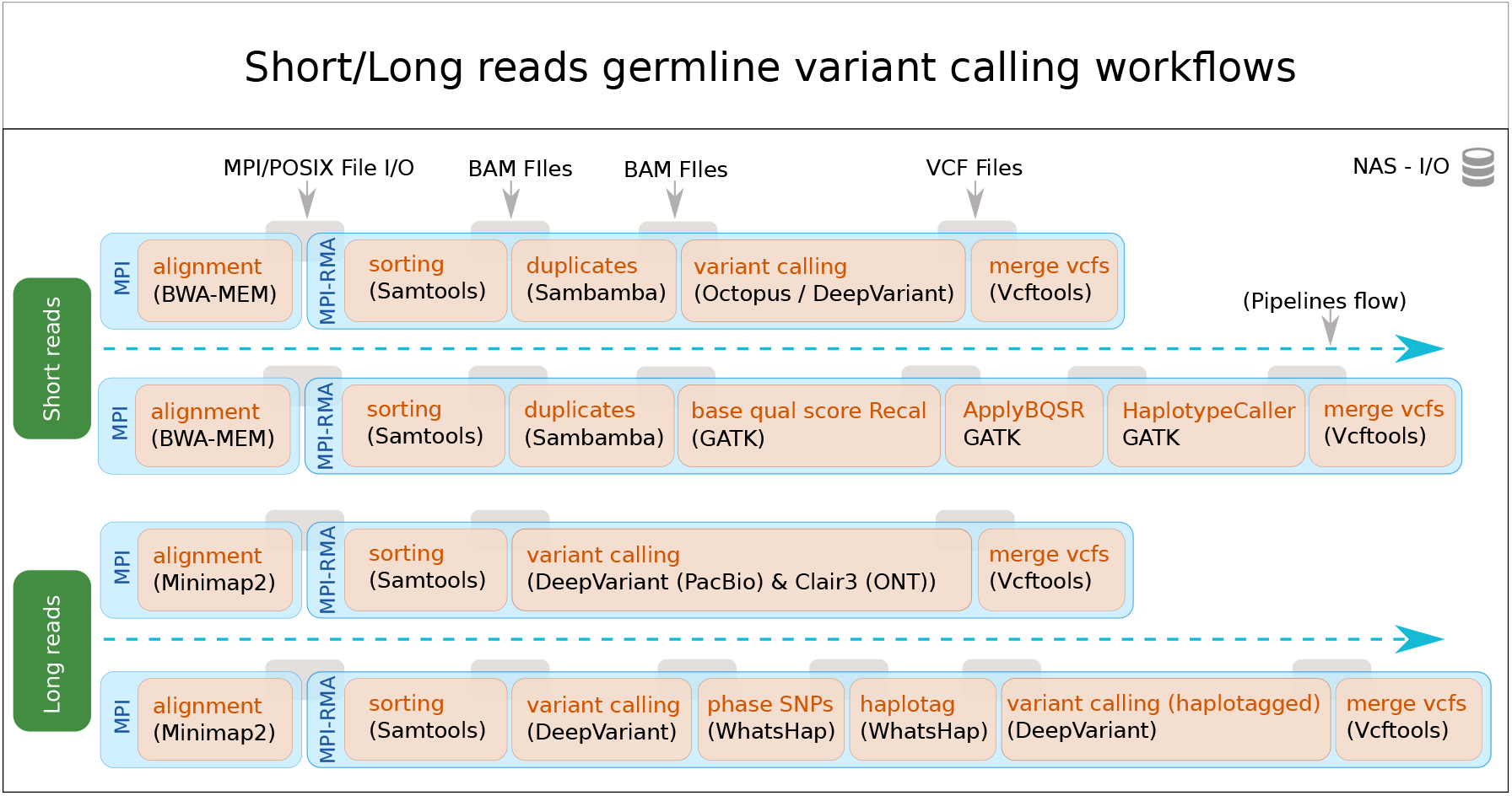
Architectural description of short and long reads based NGS data variant calling workflows.

### 2.1 Short-reads variant calling workflow

In the following subsections, we describe in details how MPI has been integrated with the BWA-MEM short read aligner and how different pre-processing applications are used in the variant calling workflow.

#### 2.1.1 Alignment

The complete algorithmic implementation details of integrating the alignment applications is given in pseudo-code representation in Algorithms 1 and 2. Normally, pair-end short reads are used for variant detection, consisting of two FASTQ raw NGS data files. As shown in Algorithm 1, we start MPI after basic parameter initialization. First, we check (Lines 5–10) if the user enabled chromosomes-based output into separate SAM files. By default this option is disabled and only single output SAM file generation is enabled. For compressed FASTQ input files (Lines 13–16), each BWA-MEM process expects equally chunked FASTQ pairs in a directory “parts” in the path of the original FASTQ files. This is done through an extra FASTQ streaming process. For streaming purpose, the user has to start an additional process; we used SeqKit Shen et al. (2016) for this purpose. SeqKit is an efficient multi-threaded command line FASTQ/FASTA data manipulation tool. In the case of uncompressed FASTQ files (Lines 18–21), we used gzseek() function which sets the file position to a given offset. We get this offset by dividing the total FASTQ file size to the number of total MPI processes. Through this technique the kseq read() function may encounter a broken first read, which we simply discard because process *N −* 1 will read the last read even if it reaches a break point for total number of bytes it can read, as shown in Algorithm 2 (Lines 1–8) in *process()* function *step-0*. In this way, we ensure that none of the input reads is dropped by any process. We calculate the size of each read and increment the bytes counter until it reaches the required number of bytes for each BWA-MEM process. Afterwards, BWA-MEM starts processing these sequences (Lines 9) in *process()* function *step-1*. Finally, we distinguish the chromosomes id for each read if writing to individual chromosome files is enabled (Lines 10–17) in *process()* function *step-2*. The output files should be written on a parallel file system with either POSIX or shared MPI I/O, depending on the best possible performance scenarios for the user.

##### Algorithm 1

Part-1: MPI integration into **BWA-MEM** and **Minimap2** for reading compressed and uncompressed FASTQ input files.

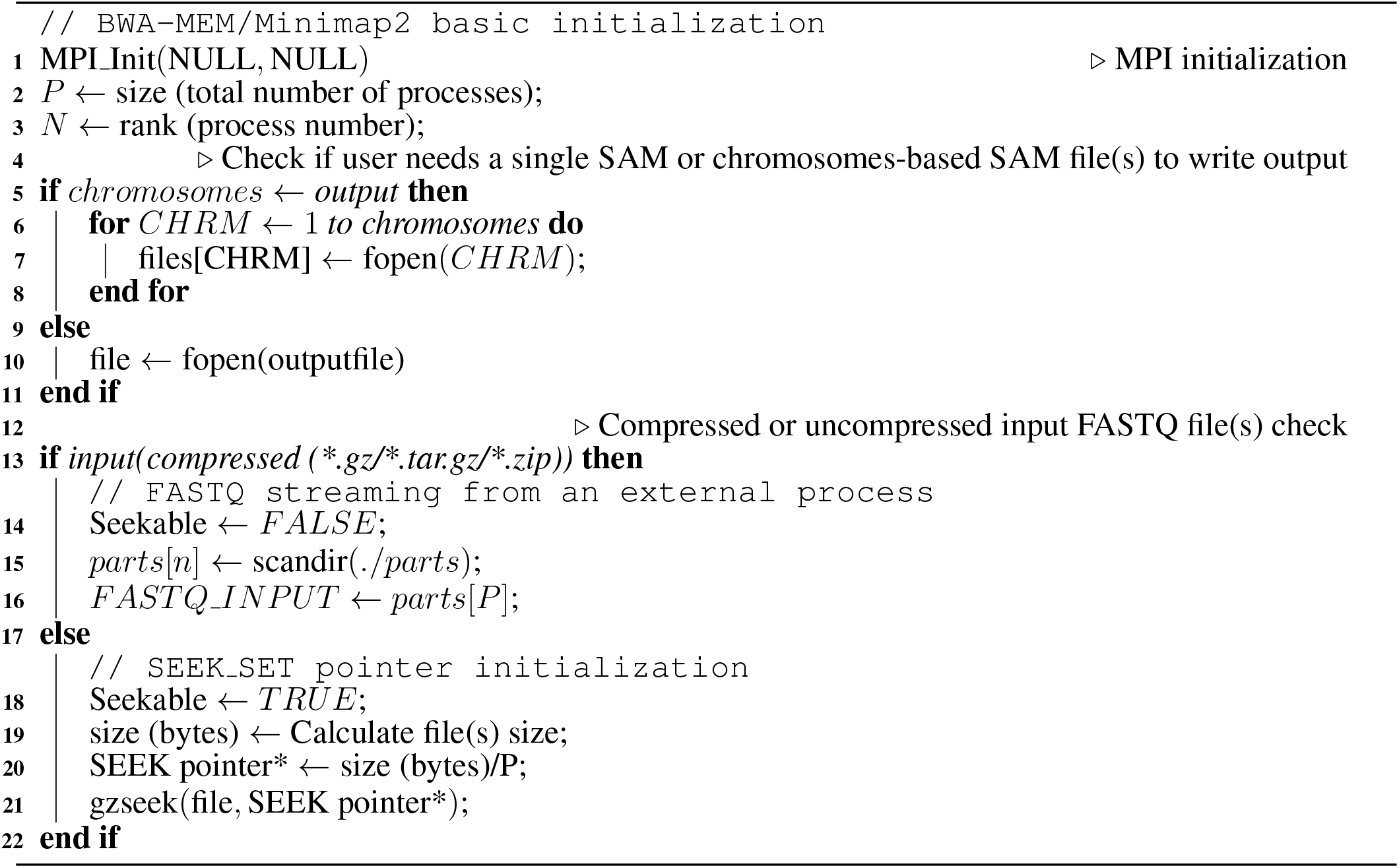

#### 2.1.2 MPI RMA-based chrosome queue

The MPI RMA interface provides applications with access to advanced features of modern high-performance networks, including direct access to remote memory through puts, gets, and atomic operations Hoefler et al. (2015). We utilize the atomic fetch-and-add functionality of MPI RMA to assign chromosomes to processes. Processes continuously increment the chromosome counter atomically until the returned value is equal to or larger than the number of chromosomes. Algorithm 3 shows the algorithm for allocating a suitable window and then querying chromosome numbers and processing them until all chromosomes have been assigned to a process. The respective window is allocated using the `“osc rdma acc single intrinsic”` info key set to true (Lines 1–2), which is supported by Open MPI and allows the implementation to utilize low-level hardware atomic operations by signaling the use of a single data element (Schuchart et al., 2019). Processes continuously increment a variable using an atomic fetch-and-op in the window to acquire the next chromosome to process (Lines 5–7) and stop once the counter exceeds the number of chromosomes available (Line 9).

##### Algorithm 2

Part-2: MPI integration into **BWA-MEM** and **Minimap2** for processing FASTQ data and saving output to SAM single or chromosome-based file.

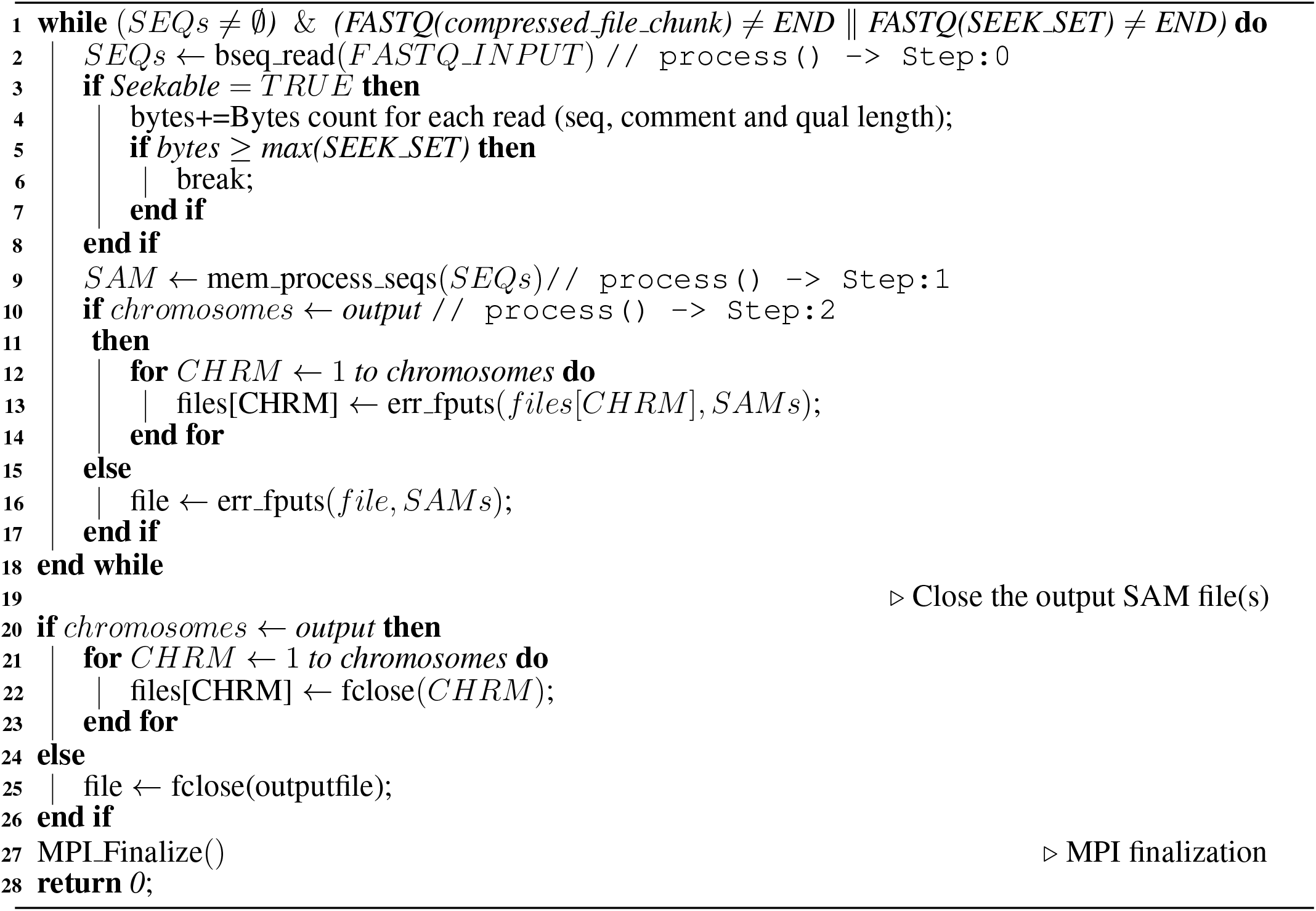

#### 2.1.3 Sorting, duplicates removal, indexing and variant calling

As described above, the MPI atomic operations based queue algorithm continuously pools the input chromosomes SAM files and operates on them for sorting, mark duplicate, index generation and variant calling algorithms. We used and integrated the sorting algorithm from Samtools and the mark duplicate algorithm from Sambamba because both perform better and have multi-threading capabilities. For variant calling, we use three different tools (GATK HaplotypeCaller, DeepVariant and Octopus) for germline variant calling of short reads. As discussed above, generating SAM files for individual chromosomes in the alignment process is not necessary, since we can also use a single SAM file. If the user wants to use all other pre-processing (sorting, mark duplicate, BAM indexing) and variant calling tools in this workflow, it is also possible to parallelize these by providing the contigs/chromosome numbers, even in the case of using a single SAM file. Finally, the individual VCFs created are merged through Bcftools to produce a final complete VCF file for further downstream analysis.

##### Algorithm 3

MPI RMA atomic operations for creating a chromosome queue for pre-procssing and variant calling

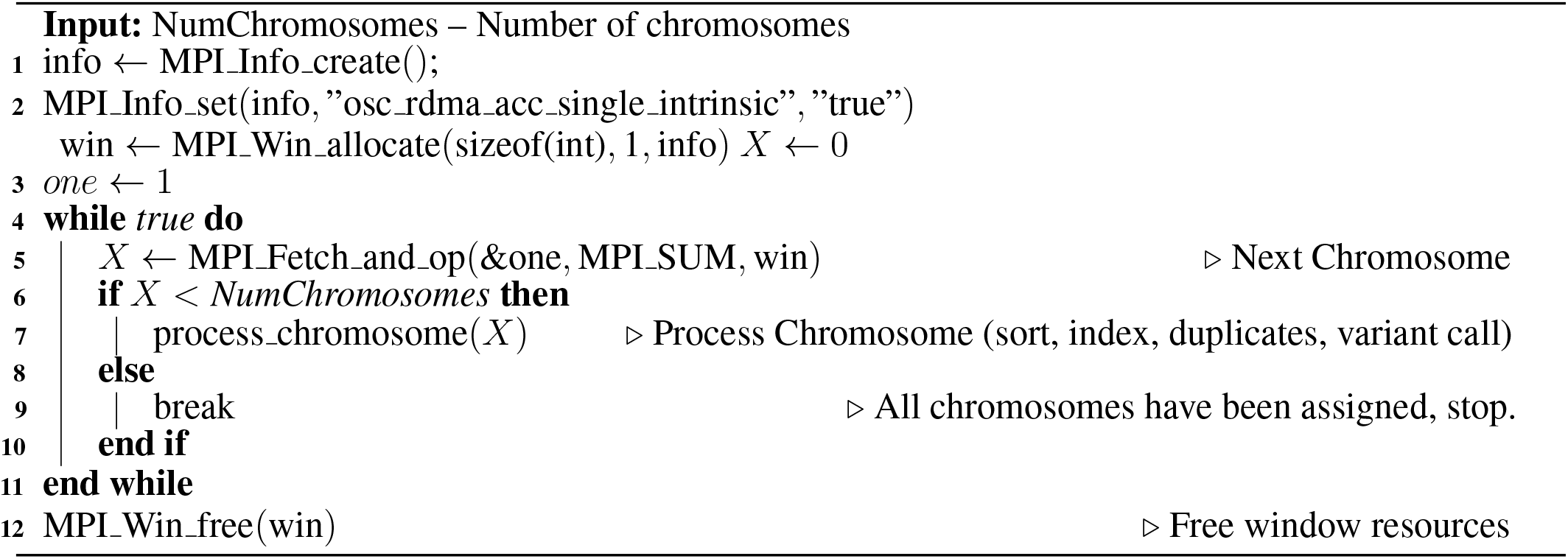

### 2.2 Long-reads variant calling workflow

In the following subsections, we describe in detail how MPI has been integrated with the long read aligner Minimap2, and how different pre-processing applications are used in the variant calling workflow.

#### 2.2.1 Alignment

Circular consensus sequencing (CCS) based PacBio HiFi reads and Oxford Nanopore technologies (ONT) long reads are making their way from research to clinical applications. They provide more in-depth and better consensus particularly in more complex repetitive regions of the genome (Wenger et al., 2019). The Minimap2 long reads aligner (with or without some additional parameter settings) is mainly being used for both PacBio and ONT long reads data. The complete algorithmic implementation details of this integration are given in pseudo-code in Algorithms 1 and 2, the detail description of Minimap2 implementation is similar to BWA-MEM as described in Section 2.1.1.

#### 2.2.2 Sorting, indexing and variant calling

For long reads after alignment, only sorting and index generation for BAM is necessary. MPI atomic operations based queue algorithm continuously polls the input chromosomes SAM files, or otherwise contigs/chromosomes number can also be used in case a single SAM file is generated through Minimap2. Then the queue algorithm operates on those files/contigs/chromosomes for sorting, BAM index generation and variant calling algorithms. We used and integrated the sorting algorithm from Samtools. For variant calling, we used three algorithms for germline variant calling of long reads (DeepVariant, DeepVariant with WhatsHap for phasing on PacBio dataset, and Clair3 for ONT dataset). These variant callers are commonly recommended by the corresponding sequencing vendors. Finally, the individual VCFs created are merged using Bcftools to produce a final complete VCF file for further downstream analysis.

## 3 RESULTS

### 3.1 Evaluation cluster

We used both HLRS Hawk (Hawk, 2021) (an HPE Apollo 9000 system at the High Performance Computing Center Stuttgart (HLRS) in Germany) and the SurfSara Snellius (Snellius, 2021) (part of the Dutch national supercomputing infrastructure) HPC clusters. Each compute node of Hawk is equipped with a dual socket AMD EPYC 7742 processors (64 cores/socket) running at 2.25 GHz. All nodes are connected through Mellanox HDR200 (interconnect bandwidth 200 Gbit/s) Infiniband adapter. Likewise, on SurfSara Snellius, each compute node is equipped with a dual socket AMD EPYC 7H12 (64 cores/socket) processors running at 2.6 GHz. All nodes are connected through Mellanox HDR100 (interconnect bandwidth 100 Gbit/s) Infiniband adapter. A local storage of 1-TBytes and the same amount of network attached storage is available on both systems.

The parallel file system available on HLRS Hawk is based on Lustre while SurfSara Snellius is equipped with IBM Spectrum Scale (GPFS) (IBM, 2020). The SLURM Workload Manager is installed on SurfSara Snellius while HLRS Hawk uses PBSPro workload manager and job scheduler.

### 3.2 Datasets

We used the Illumina, PacBio HiFi, and ONT HG002 datasets taken from the PrecisionFDA challenge V2 (FDA, 2019)) dataset for variant calling workflows. We also used 300x sequencing coverage WGS data from Genome in a Bottle (GIAB) aligned with novoalign for the Illumina HiSeq 300x reads for NA12878 GIAB (2020) to analyze the scalability of BWA-MEM aligners. Human genome reference GRCh38 (UCSC, 2019) is used as a reference genome. For accuracy comparisons, the GIAB v4.2.1 (GIAB, 2021) benchmark set for HG002 dataset is used.

### 3.3 Runtime performance

In this section, we analyze and compare the runtime performance of MPI based aligners (BWA-MEM and Minimap2) with existing state-of-the-art MPI and Apache Spark based implementations. We also benchmark the performance of variant calling workflows using different variant callers on a cluster of up to 8 nodes.

#### 3.3.1 Short reads alignment

We compare the scalability and runtime performance of GenMPI with those based on Apache Spark (ADAM’s Cannoli (Massie et al., 2013) alignment) and another MPI-based implementation, QUARTIC (Frédéric et al., 2020) (mpiBWA). We use different numbers of nodes; 2, 4, 8 and 16 nodes for runtime performance comparisons. ADAM’s Cannoli uses the built-in Scala API from the Apache Spark backend for distributing and scheduling data for parallel processing. The Cannoli wrapper encompasses many different aligners and variant callers for distributed processing. Apache Spark based implementations use HDFS or NFS for I/O operations. QUARTIC (mpiBWA) is a distributed BWA-MEM alignment algorithm employing MPI functionality and uses MPI shared I/O for input/output on parallel file system. In Figure 2 (left bar), we show the BWA-MEM runtime on a single node utilizing all 128 system CPU cores with one thread for each core. We consider this time as an ideal runtime for a single node. When we compare this runtime by increasing the number of nodes in a cluster as shown in Figure 2, we see QUARTIC and GenMPI perform better than this ideal runtime, while ADAM Cannoli BWA-MEM implementation has less than ideal scalability. We attribute the super-linear scaling of both QUARTIC and GenMPI to scalability issues of BWA-MEM when utilizing all cores on a single node using a single process (128 threads). As a consequence, we use two MPI processes on each node to overcome this inefficiency for all other runs. Overall GenMPI outperforms QUARTIC by almost 20% in terms of runtime by reducing read/write I/O time, and outperforms Apache Spark based ADAM Cannoli BWA-MEM by 2x in terms of runtime.

**Figure 2.**
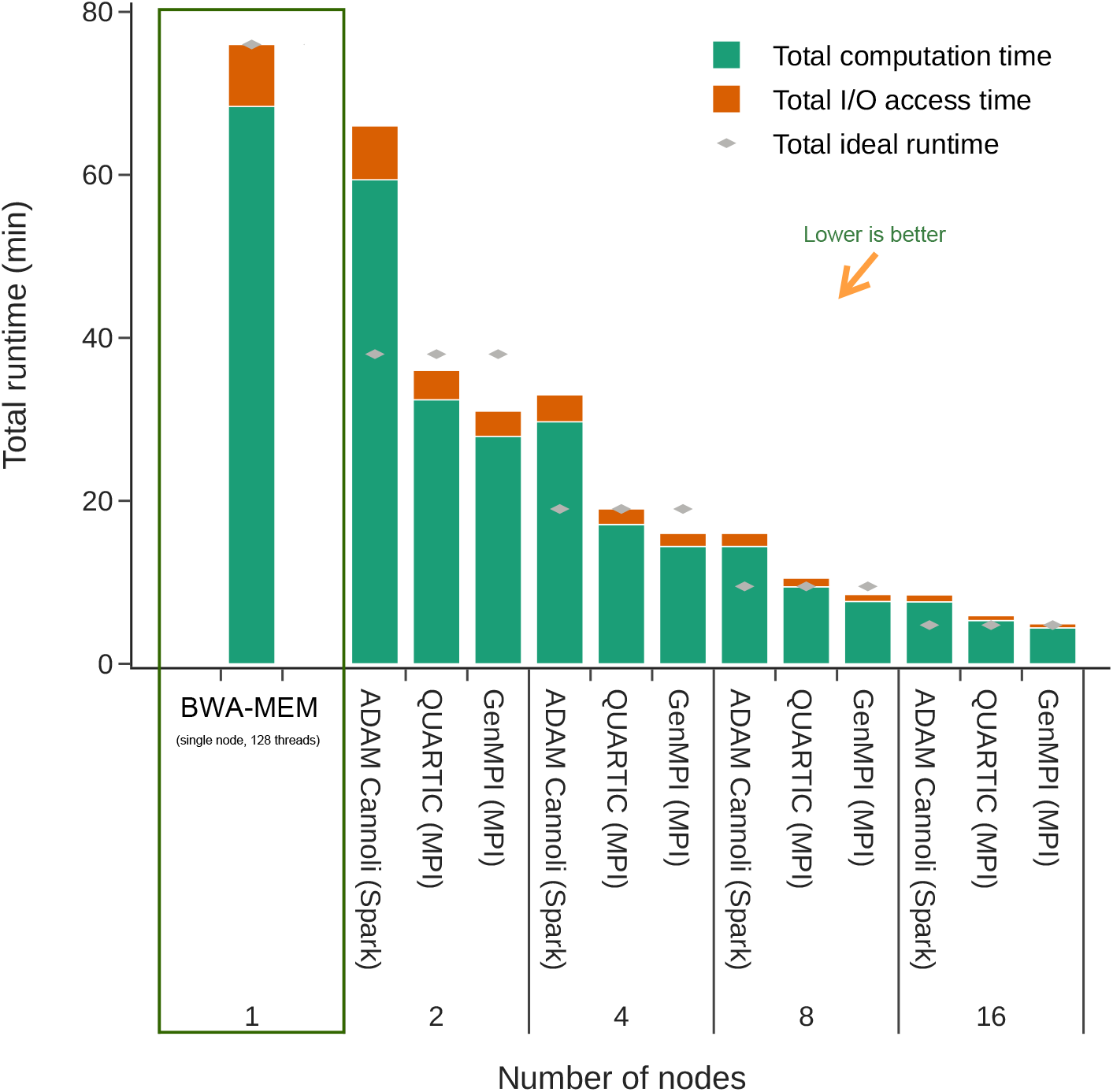
Run-time and scalability comparisons of ADAM’s Cannoli, QUARTIC mpiBWA and this work implementations of BWA-MEM aligner on a cluster of varying number of nodes using HG002 (NA24385) Illumina NovaSeq 35x coverage dataset. Ideal runtime is the runtime assuming ideal scalability on the available nodes.

#### 3.3.2 Short-reads variant calling workflow

As discussed in the alignment section, we store BWA-MEM SAM output in chromosomes files. Sorting can be performed using existing tools on these chromosome files in parallel on the cluster. We use *Samtools’* sorting algorithm since it is one of the most efficient due to its in-memory and multi-threading functionality. Similarly, for duplicate removal we use *Picard*’s MarkDuplicate compatible algorithm *Sambamba*, which is also multi-threaded. We also use GATK best practices pipeline applications like *Base quality score recalibration* (BQSR), *ApplyBQSR*, and *HaplotypeCaller* afterwards. Moreover, we integrate *DeepVariant* and *Octopus*, both recent and accurate variant callers, in this workflow. Their published results show high accuracy and *F*_1_-score compared to other state-of-the-art variant callers like *GATK4 HaplotypeCaller* (Institute, 2010), *Strelka2* (Kim et al., 2018), or *FreeBayes* (Garrison and Marth, 2012).

We compare the total runtime of these workflows on varying numbers of nodes. As shown in Figure 3, moving from a single node to two nodes a more than 3x runtime speedup is achieved because we allocated multiple MPI processes for pre-processing and variant calling applications on each node as these tools have some single node (128 CPU cores) scalability limitations. Increasing the cluster size from 4 to 8 nodes yields only 70-80% runtime improvements because of poor chromosome load-balance. The same runtime speedup is observed for DeepVariant instead of Octopus. On the other hand, GATK best practices pipeline applications are either slow or single-threaded; therefore we only focus on optimizing the workflow by insuring accuracy while running with maximum efficiency on a single node. As shown in Figure 4, using alternative application like Samtools/Sambamba instead of Picard (sort and markdup) respectively, and parallel execution of GATK BQSR, ApplyBQSR and HaplotypeCaller applications for chromosomes we have achieved more than 6.5x runtime speedup compared to the baseline. Since GATK processes chromosomes using a single thread, distributing it’s execution across multiple nodes is not worthwhile due to the limited number of chromosomes. We have only 25 chromosomes which can be ran in parallel on a single node more efficiently.

**Figure 3.**
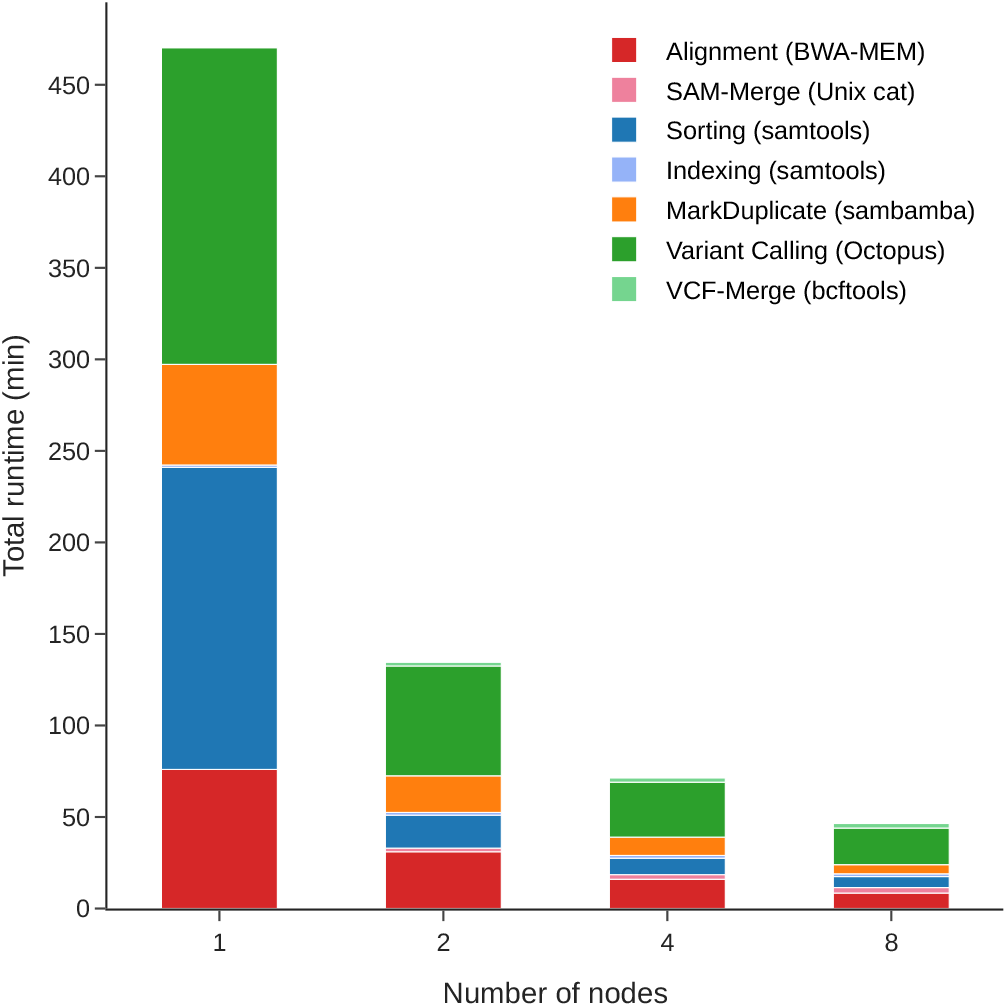
Run-time and scalability benchmark of GenMPI for BWA-MEM aligner and Octopus variant caller on a cluster of varying number of nodes using HG002 (NA24385) Illumina NovaSeq 35x coverage dataset.

**Figure 4.**
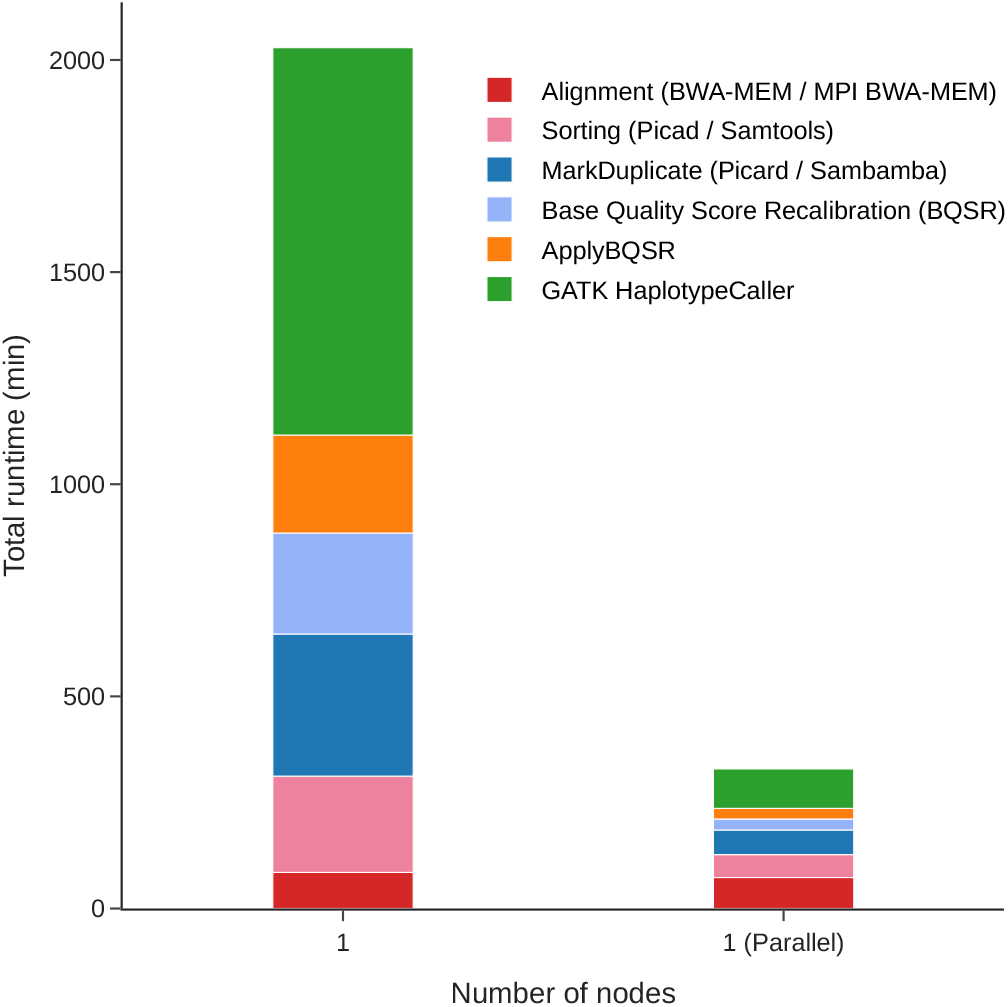
Run-time and scalability benchmark of GenMPI for BWA-MEM aligner and GATK best practices pipeline on a single node versus alternative parallel single node methods using HG002 (NA24385) Illumina NovaSeq 35x coverage dataset.

#### 3.3.3 Long reads alignment

GenMPI is the first-ever cluster scale implementation of any long reads aligners. As discussed in Section 2, both chromosomes based SAM files and a single SAM file output options are implemented in Minimap2. Figure 5 shows the total runtime for long-reads aligner, Miniamap2, which exhibits close to linear scalability when increasing the number of nodes in the cluster. The ideal theoretical values for linear scalability are also shown. As shown in the figure, there is a minor performance degradation compared to ideal runtimes that is caused by read/write I/O operations for long reads. Due to the efficient performance of network-attached parallel file system, the overhead of storing SAM data in multiple chromosomes file is minimal compared to a single SAM file generation from all the MPI processes.

**Figure 5.**
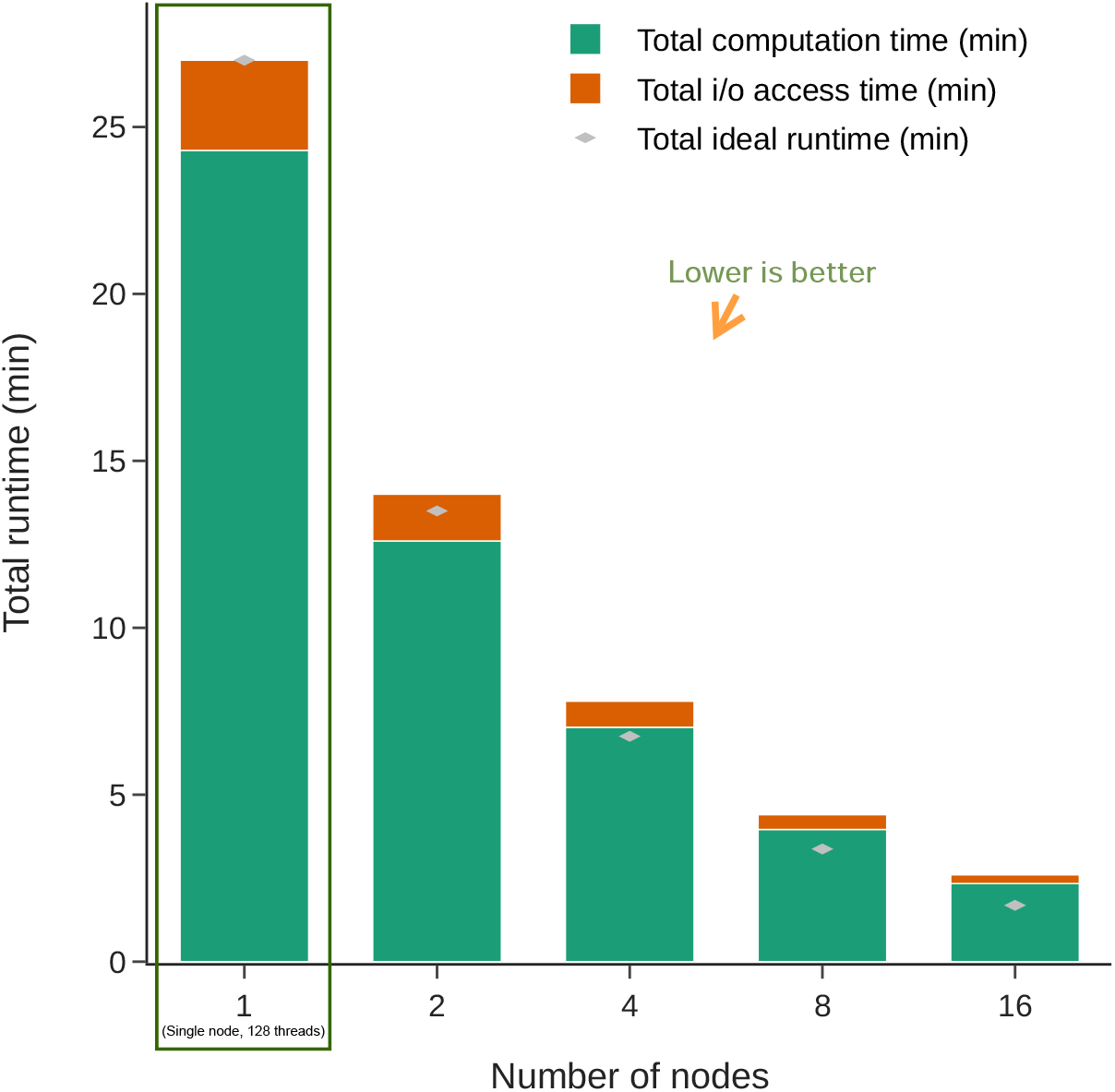
Run-time and scalability performance of MPI based Minimap2 implementation. HG002(NA24385) PacBio HiFi 35x coverage dataset has been used on a cluster of varying number of nodes.

#### 3.3.4 Long reads variant calling workflow

Almost all new long reads variant callers only require aligned and sorted reads. Long reads sorting is performed through Samtools ‘sort’ algorithm. As mentioned before, for PacBio data, we used DeepVariant and DeepVariant with WhatsHap (Martin et al., 2016). WhatsHap is used to reconstruct the chromosomes haplotypes and then write out the input VCF augmented with phasing information in its first pass, in the second pass WhatsHap haplotag writes information of reads along with the variants. This information can be used again in DeepVariant for better INDEL detection and accuracy purposes. In Figure 6, we show the scalability results of long reads variant calling workflow (Minimap2, Samtools sorting and DeepVariant) on a cluster where more than 2.25x runtime speedup is achieved from 1 node to 2 node cluster and a total of 5x speedup is achieved on a 8 node cluster. This scalability is again only valid up to 8 nodes since we aim at reproducing the exact same variant output as that of a single node, which means that we do not sub-divide chromosomes into smaller pieces. Similarly, for ONT dataset we used Clair3 which provides better accuracy and performance for both SNP and INDEL variants as compared to other variant callers on ONT data. As shown in Figure 7 due to some internal scalability limitations of Clair3 we are only able to achieve 1.5x runtime speedup for two nodes cluster as compared to a single node runtime and similar limited scalability trend is shown for more nodes.

**Figure 6.**
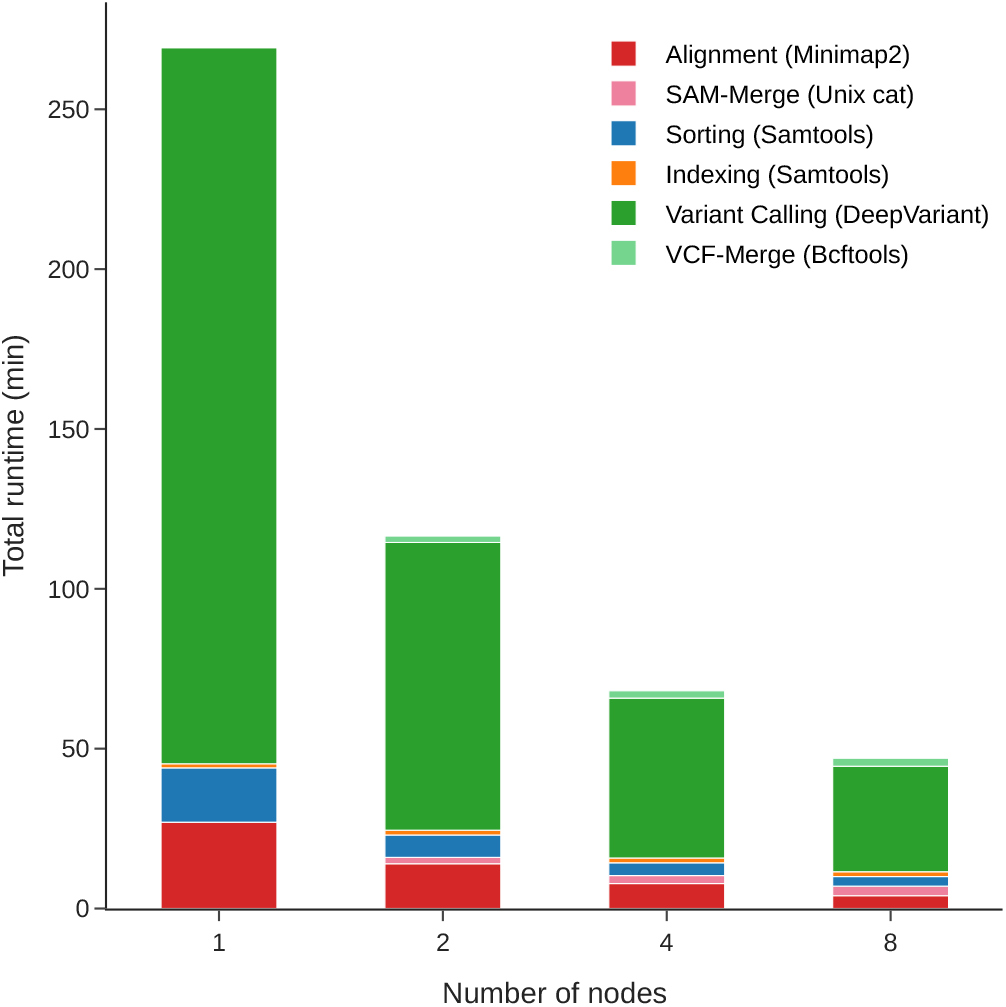
Run-time and scalability benchmark of this MPI based implementation for Minimap2 aligner and DeepVariant variant caller on a cluster of varying number of nodes using HG002 (NA24385) PacBio HiFi 35x coverage dataset.

**Figure 7.**
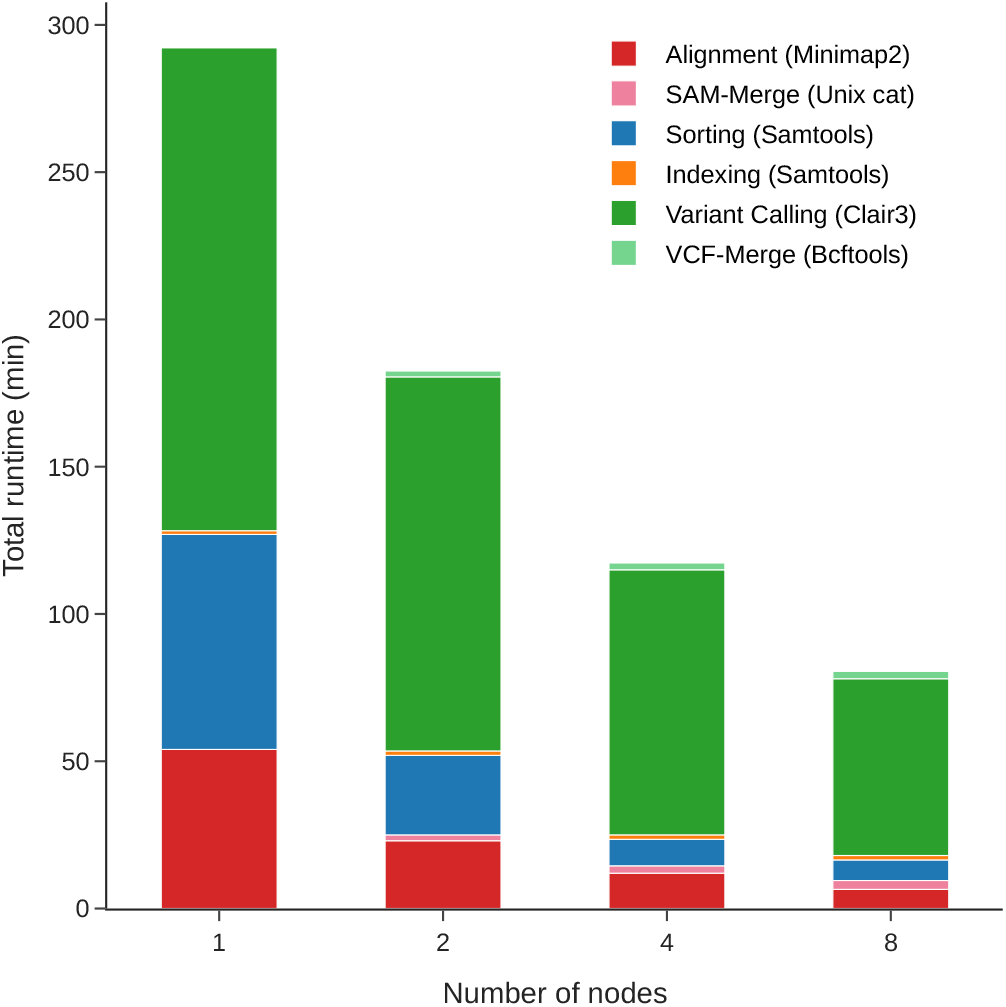
Run-time and scalability benchmark of this MPI based implementation for Minimap2 aligner and Clair3 variant caller on a cluster of varying number of nodes using HG002 (NA24385) ONT Guppy 3.6.0 dataset.

### 3.4 Accuracy

In this subsection, we present small variants (SNP and INDEL) detection accuracy. The GA4GH small variant benchmarking tool hap.py has been used to compare the variants in all methods. We compared all these statistics on Chr1-22 and X, Y for each dataset. We compared the small variants (SNP and INDEL) detection accuracy for both short and long reads on HG002 (PrecisionFDA challenge V2) dataset against GIAB v4.2.1 benchmark set. We only tested the accuracy metrics for all-benchmarking region of GIAB v4.2.1 benchmark.

#### 3.4.1 SNP accuracy

Figure 8 shows the accuracy performance of SNPs in terms of precision and recall for short reads methods (GATK HaplotypeCaller, Octopus and DeepVariant) on Illumina dataset and long reads methods (DeepVariant, DeepVariant with WhatsHap and Clair3) on PacBio and ONT datasets respectively. Both DeepVariant and DeepVariant with WhatsHap on PacBio data perform best as compared to other methods while GATK HaplotypeCaller SNP performance is well below all other methods. Overall, the best SNP *F*_1_ score is 0.999283 for PacBio dataset using DeepVariant variant caller, as shown in Table 2.

**Figure 8.**
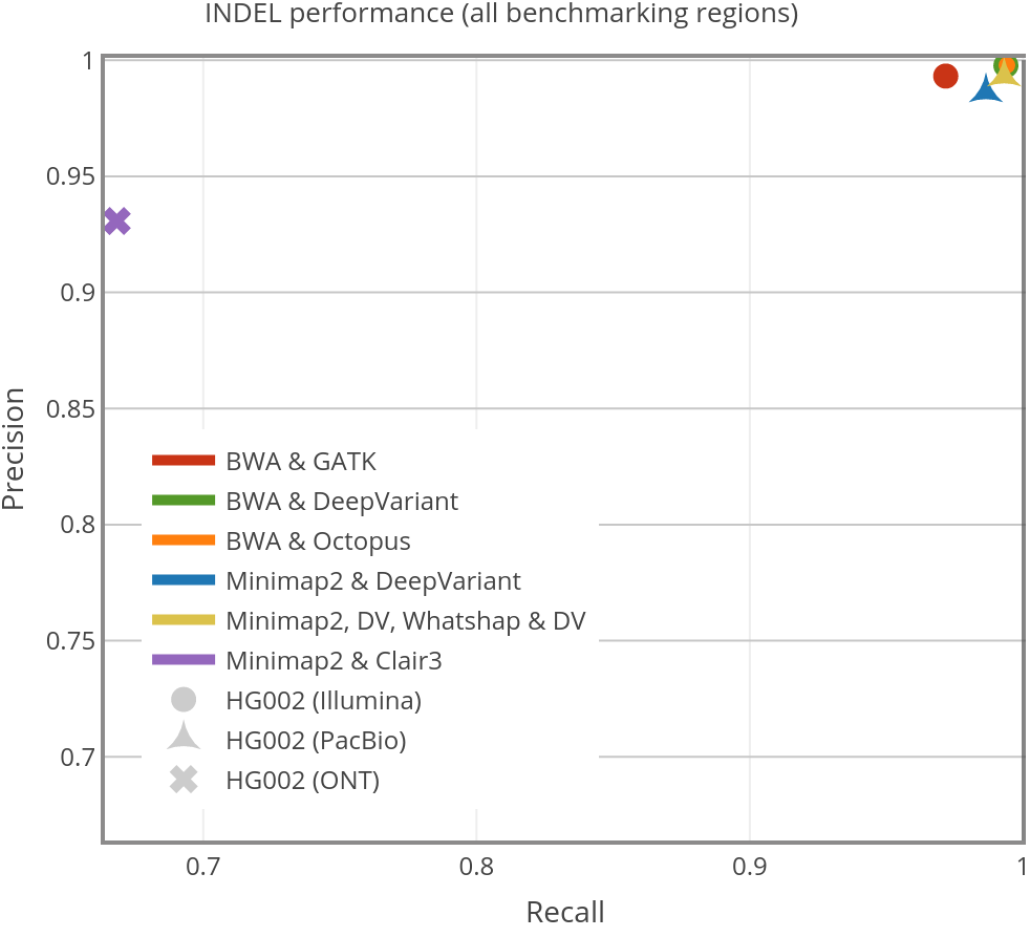
INDEL performance comparison of different short and long reads variant calling methods on HG002 dataset.

#### 3.4.2 INDELs accuracy

Similarly in Figure 9 the accuracy performance in terms of precision and recall of INDELs for short reads methods (GATK HaplotypeCaller, Octopus and DeepVariant) on Illumina dataset and long reads methods (DeepVariant, DeepVariant with WhatsHap and Clair3) on PacBio and ONT datasets respectively has been shown. We have observed for INDEL performance that Illumina short read dataset seems to performs best as compared to other datasets for both DeepVariant and Octopus methods. ONT dataset INDEL performance is lower than any other datasets. Overall the best INDEL *F*_1_ score is 0.995957 for Illumina dataset using Octopus variant caller as shown in Table 1.

**Table 1.**
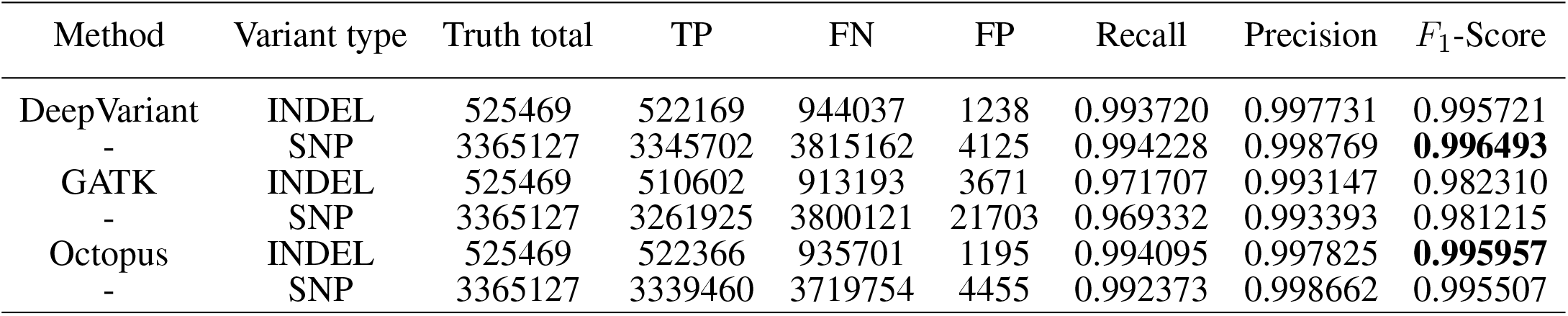
Accuracy evaluation of small variants for HG002 (pair-end Illumina short reads) against GIAB HG002 v4.2 benchmarking set for different methods adopted in this workflow. This table shows the SNP <u>and INDEL results for all benchmarking regions.</u>

| Method | Variant type | Truth total | TP | FN | FP | Recall | Precision | $F_1$-Score |
| --- | --- | --- | --- | --- | --- | --- | --- | --- |
| DeepVariant | INDEL | 525469 | 522169 | 944037 | 1238 | 0.993720 | 0.997731 | 0.995721 |
| - | SNP | 3365127 | 3345702 | 3815162 | 4125 | 0.994228 | 0.998769 | **0.996493** |
| GATK | INDEL | 525469 | 510602 | 913193 | 3671 | 0.971707 | 0.993147 | 0.982310 |
| - | SNP | 3365127 | 3261925 | 3800121 | 21703 | 0.969332 | 0.993393 | 0.981215 |
| Octopus | INDEL | 525469 | 522366 | 935701 | 1195 | 0.994095 | 0.997825 | **0.995957** |
| - | SNP | 3365127 | 3339460 | 3719754 | 4455 | 0.992373 | 0.998662 | 0.995507 |

**Figure 9.**
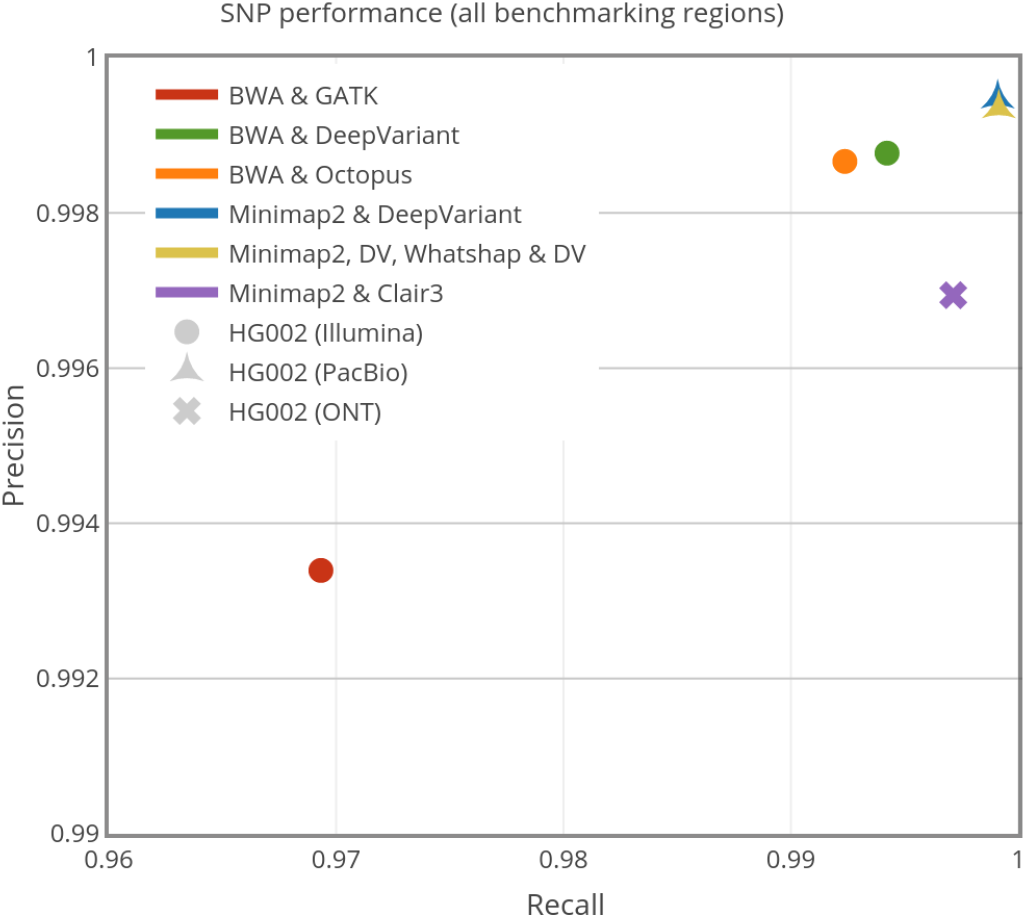
SNP performance comparison of different short and long reads variant calling methods on HG002 dataset.

## 4 DISCUSSION

This section further elaborates on the benefits of this approach in a broader context of its applicability in real-time usage on HPC clusters.

### 4.1 Runtimes

Total runtimes of the used aligners show a good linear scalability on small to big clusters in both single SAM output as well as for chromosomes based SAM output. Similarly, for complete variant calling workflows almost all methods provide more than 2x to 6x lower runtime when executed on 2 to 8 cluster nodes, respectively. Because GATK best practices pipeline applications are not multi-threaded, cluster scalability is not possible. Neverthelesse, we have achieved more than 6.5x speedup when executing in parallel on a single node. Similarly, Clair3 has multi-threading limitations that preclude proper scaling across multiple nodes of a cluster.

### 4.2 Accuracy and reproducibility

Accuracy results for the Illumina short-read dataset using GATK HaplotypeCaller, Octopus, and DeepVariant variant callers are shown in Table 1. Similarly, Table 2 shows accuracy results for the PacBio dataset using the DeepVariant and DeepVariant with WhatsHap variant callers. Table 3 lists the accuracy results for the ONT dataset using the Clair3 variant caller. These results are obtained through GA4GH small variant benchmarking tool hap.py.

**Table 2.** Accuracy evaluation of small variants for HG002 (PacBio) against GIAB HG002 v4.2 benchmarking set for different methods adopted in this workflow. This table shows the SNP and INDEL <u>results for all benchmarking regions.</u>

| Method | Variant type | Truth total | TP | FN | FP | Recall | Precision | $F_1$-Score |
| --- | --- | --- | --- | --- | --- | --- | --- | --- |
| DeepVariant | INDEL | 525469 | 518336 | 970119 | 7560 | 0.986425 | 0.986187 | 0.986306 |
| - | SNP | 3365127 | 3362077 | 4054200 | 1778 | 0.999094 | 0.999472 | **0.999283** |
| DV-WH-DV | INDEL | 525469 | 521828 | 981392 | 3844 | 0.993071 | 0.992977 | **0.993024** |
| - | SNP | 3365127 | 3362239 | 4060995 | 2199 | 0.999142 | 0.999347 | 0.999244 |

**Table 3.** Accuracy evaluation of small variants for HG002 (ONT) against GIAB HG002 v4.2 benchmarking set for different methods adopted in this workflow. This table shows the SNP and INDEL results for all <u>benchmarking regions.</u>

| Method | Variant type | Truth total | TP | FN | FP | Recall | Precision | $F_1$-Score |
| --- | --- | --- | --- | --- | --- | --- | --- | --- |
| Clair3 | INDEL | 525469 | 351200 | 554604 | 26874 | 0.668355 | 0.930726 | **0.778016** |
| - | SNP | 3365127 | 3355502 | 4148864 | 10306 | 0.997140 | 0.996939 | **0.997039** |

The accuracy comparisons between the MPI versions of the pipelines and their single node baseline show that these workflows produce the exact same recall, precision, and *F*_1_-score for both short and long reads based variant calling workflows. This ensures the reproducibility of original BWA-MEM and Minimap2 functionality when using GenMPI.

### 4.3 Scalability

The MPI parallelization of both aligners (BWA-MEM and Minimap2) is highly scalable. To further evaluate the scalability of both aligners, we tested the GIAB 300x coverage whole genome sample NA12878 for HG001 with the GenMPI using BWA-MEM. Figure 10 shows scalability results of aligning almost 2.5 TBytes data in just 10 minutes walltime on a 64 nodes. We show both computation and I/O time for both aligners separately. In both aligners, the resultant graphs show that increasing the number of nodes in the cluster, the runtime of alignments decreases almost linearly. We also observed that I/O time is increasing slightly when increasing the number of nodes but it is not a bottleneck for the scalability.

**Figure 10.**
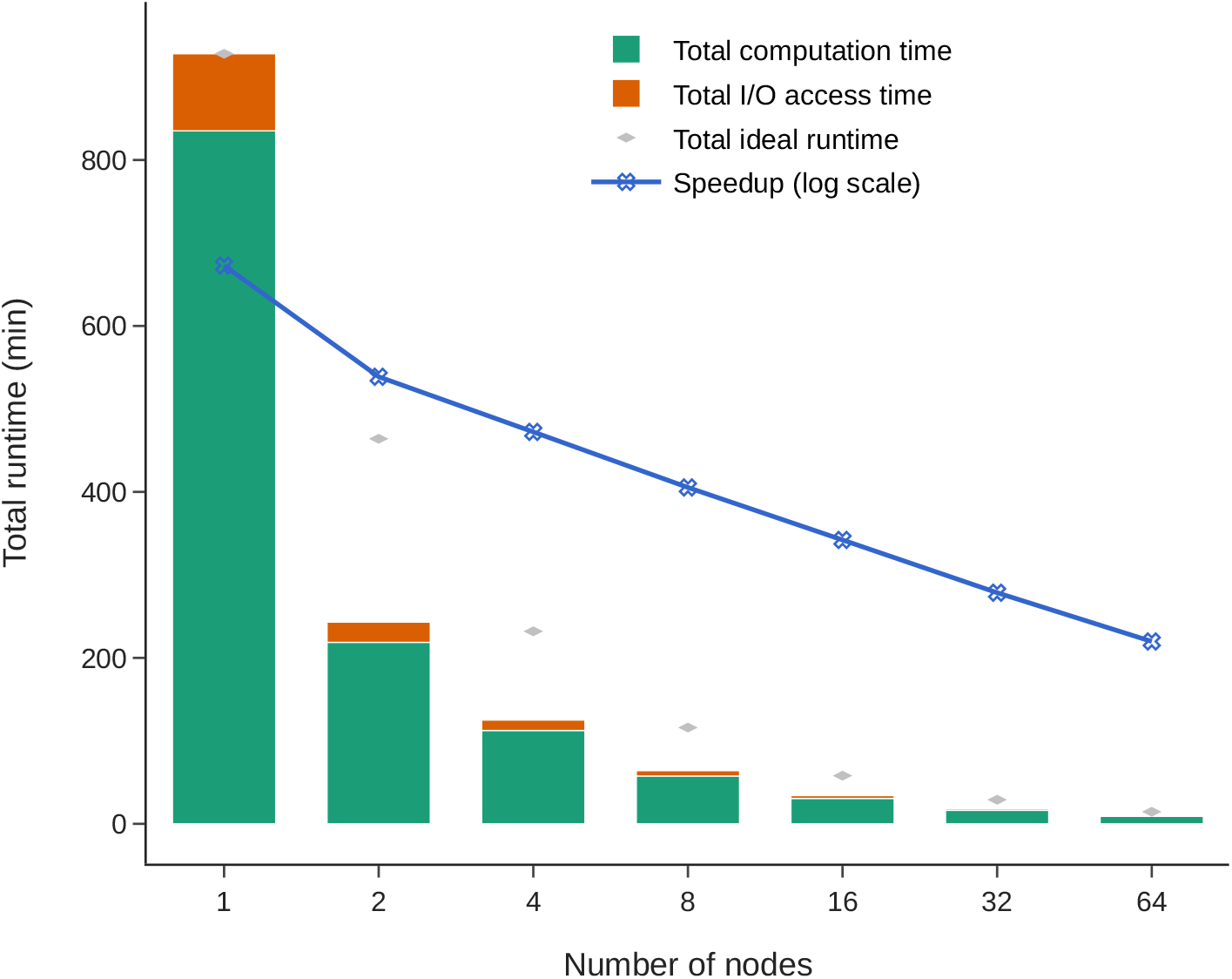
GenMPI BWA-MEM performance and scalability measurements for 300x sequencing coverage WGS data from Genome in a Bottle (GIAB) aligned with novoalign for the Illumina HiSeq 300x reads for NA12878 GIAB (2020).

The only issue is in shared MPI I/O, particularly in writing and reading part of Minimap2 for long reads is a limiting factor for linear scalability. Both MPI I/O and POSIX file system options to output a single SAM file, chromosomes-based SAM output and chromosomes-regions specific SAM output have been implemented as shown in Figure 11. We have observed significant overhead due to synchronization effects in writing to POSIX I/O as well as MPI I/O when a single output SAM file is written, as shown in Figure 11(a). For per-chromosome SAM output (Figure 11(b)), the best suitable option is using POSIX I/O. For the chromosomes-regions specific SAM output option (Figure 11(c)), we used 128 output files, which provides comparable results to per-chromosome SAM output with POSIX I/O but still affects the overall performance compared to ideal runtime. The reason behind this is inner reads identification and writing loops which direct reads to a specific file. We have tested both MPI I/O blocking and non-blocking I/O operations for writing SAM output. Due to processes synchronization for wiring SAM data chunks to I/O, an extra overhead slows down the writing of results. This overhead can be mitigated by directly integrating aligners with sorting and indexing applications without relying on file I/O. This will be part of our future work (Section 4.7).

**Figure 11.**
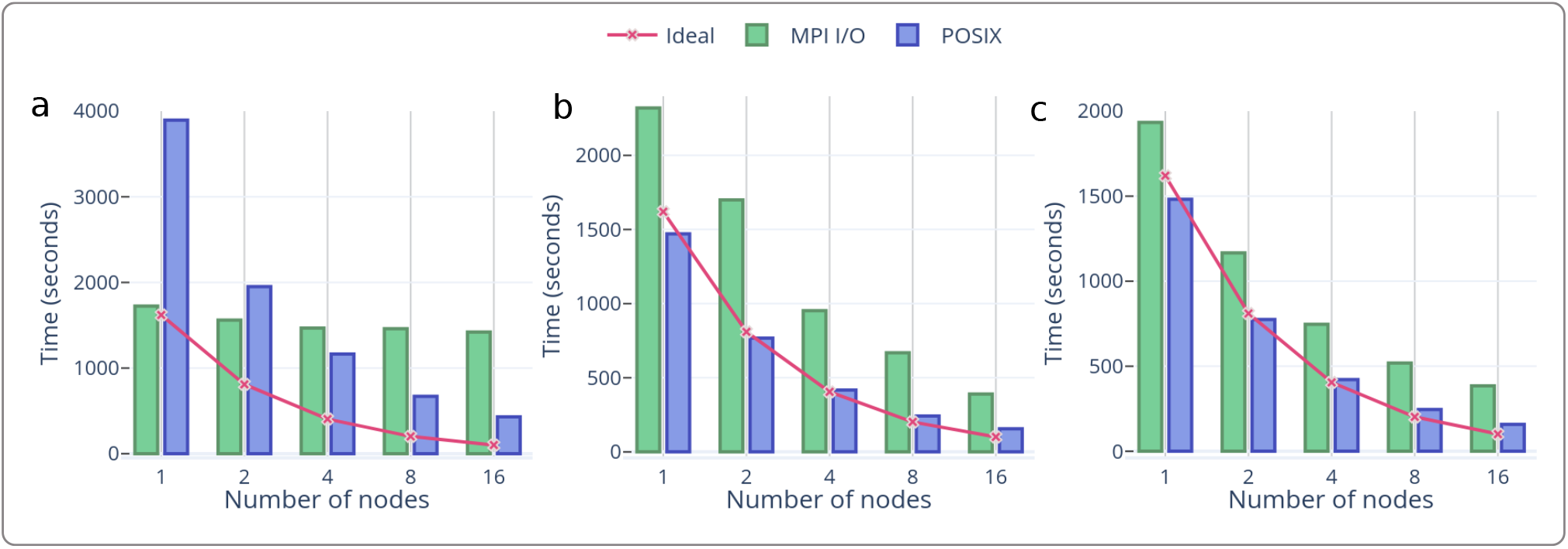
Performance and scalability comparisons when using MPI shared I/O and POSIX I/O for (a) single SAM output (b) chromosomes based SAM output and (c) chromosomes regions specific SAM output.

### 4.4 Portability and deployment

By using the MPI standard, this workflow is portable and easily deployable to any HPC cluster. A detailed description and quick start guide to run all methods in this approach are given on the project github page. We also tested this implementation with OpenMPI, Intel MPI and HPE MPI flavors and it compiles/runs appropriately.

### 4.5 Cost efficiency

Our cost estimations predict that a significant amount can be saved when opting for public clouds clusters instead of a single large node in the cloud. Particularly in BWA-MEM alignment, clusters utilize maximum system resources resulting in cost saving of more than 50%. Similarly, DeepVariant and Octopus variant callers have some limitation in single node performance for a higher number of cores per socket. More than 20-30% cost can be saved using these variant callers on a cluster with each node having two sockets each with 64 CPU cores.

### 4.6 Memory consumption

MPI based scalable implementations can have a large edge over Apache Spark based variant calling workflows in terms of memory consumption. The MPI implementations do not need extra memory for the platform itself nor to store the data during data shuffling in Spark based implementations.

### 4.7 Future work

This whole workflow implementation (GenMPI) uses storage for intermediate applications I/O read and write operations in form of SAM/BAM files. For future work, integrating pre-processing applications like GenMPI BWA-MEM with sorting (mpiSORT (Frédéric et al., 2020)) and index generation would yield further performance gains. We have observed that 5-10% of the total time in GenMPI BWA-MEM and 50% of the total time in mpiSORT are spent on file I/O, which could be mitigated through this approach.

## 5 CONCLUSION

In this work, we have implemented GenMPI, an MPI based scalable method for both widely used short and long reads aligners, BWA-MEM and Minimap2, respectively. One of the main goals of GenMPI is to ensure 100% identical variant output compared to the single node baseline. In addition, GenMPI provides a flexible architecture which can be used to integrate a variety of alignment and variant calling tools. Compressed and uncompressed FASTQ input is supported and output can be a single SAM file or chromosomes-based SAM files generated by MPI processes. This output can be stored on network attached storage through both POSIX and MPI shared I/O, whichever is convenient and efficient for the underlying HPC system. Results show that these implementations outperform existing Apache Spark based implementation of alignment algorithms by 2x and yield a 20% speedup over state-of-the-art MPI implementations of the BWA-MEM algorithm. Likewise, we also integrate pre-processing applications (sorting, indexing, duplicates removal (for short reads)) as well as variant callers like GATK HaplotypeCaller, DeepVariant, Octopus and Clair3 for both short (Illumina) and long reads (PacBio and ONT) datasets. The variant calling workflows are scalable for up to 8 nodes cluster while giving 2x to 6x total runtime speedups.

Particularly, we show that the distribution of chromosomes across aligners is almost linearly scalable, even when tested using 300x coverage datasets on up to 64 nodes cluster for BWA-MEM. At scale, the alignment completes in under 10 minutes walltime. Thanks to the use of MPI, the workflow implementation is portable and easily deployable on any public cloud or private HPC cluster with minimal effort. Memory requirements do not exceed the actual software needs. The final accuracy results (Recall, Precision, and F_1_ score) for variant calling have been reproduced with hap.py against latest GIAB benchmarks set and are shown to be identical to those of the baseline pipeline.

## 6 ADDITIONAL REQUIREMENTS

For additional requirements for specific article types and further information please refer to Author Guidelines.

## AUTHOR CONTRIBUTIONS

Z.A.A, P.H., J.G. and C.N. conceived and supervised this work. T.A. and J.S. implemented and developed MPI-based whole variant calling workflow including MPI integration into aligners. All authors read and approved the final manuscript.

## ACKNOWLEDGMENTS

We are grateful for the resources provided by the High Performance Computing Center Stuttgart (HLRS). Special thanks to Alexey Cheptsov (HLRS) for all the HPC related support. This work was also carried out in part on the Dutch national e-infrastructure with the support of SURF Cooperative.

## SUPPLEMENTAL DATA

N/A

## DATA AVAILABILITY STATEMENT

All the codes and scripts are available at: https://github.com/abs-tudelft/gen-mpi. Illumina, PacBio HIFI and ONT HG002 datasets taken from PrecisionFDA challenge V2 (FDA, 2019)). We also used 300x sequencing coverage WGS data from Genome in a Bottle (GIAB) aligned with novoalign for the Illumina HiSeq 300x reads for NA12878 GIAB (2020). Human genome reference GRCh38 (UCSC, 2019) is used as a reference genome. For accuracy comparisons, GIAB v4.2.1 (GIAB, 2021) benchmark set for HG002 dataset is used.

